# Molecular dynamics simulations of protein aggregation: protocols for simulation setup and analysis with Markov state models and transition networks

**DOI:** 10.1101/2020.04.25.060269

**Authors:** Suman Samantray, Wibke Schumann, Alexander-Maurice Illig, Martin Carballo-Pacheco, Arghadwip Paul, Bogdan Barz, Birgit Strodel

## Abstract

Protein disorder and aggregation play significant roles in the pathogenesis of numerous neurodegenerative diseases, such as Alzheimer’s and Parkinson’s disease. The end products of the aggregation process in these diseases are β-sheet rich amyloid fibrils. Though in most cases small, soluble oligomers formed during amyloid aggregation are the toxic species. A full understanding of the physicochemical forces behind the protein aggregation process is required if one aims to reveal the molecular basis of the various amyloid diseases. Among a multitude of biophysical and biochemical techniques that are employed for studying protein aggregation, molecular dynamics (MD) simulations at the atomic level provide the highest temporal and spatial resolution of this process, capturing key steps during the formation of amyloid oligomers. Here we provide a step-by-step guide for setting up, running, and analyzing MD simulations of aggregating peptides using GROMACS. For the analysis we provide the scripts that were developed in our lab, which allow to determine the oligomer size and inter-peptide contacts that drive the aggregation process. Moreover, we explain and provide the tools to derive Markov state models and transition networks from MD data of peptide aggregation.

## 1 Introduction

During protein aggregation, misfolded or intrinsically disordered proteins assemble first into oligomers, which can grow into highly-ordered β-sheet aggregates called amyloid fibrils, which, depending on the protein, takes place in the intra- or extracellular environment. This process is highly associated with various, often neurodegenerative diseases, such as Alzheimer’s and Parkinson’s diseases [1,2]. Neurodegenerative diseases are debilitating conditions that result in progressive degeneration and/or death of nerve cells, causing problems with movement (called ataxia) and/or mental functioning (called dementia). To our knowledge, none of these diseases linked to amyloid aggregation are currently curable and finding a cure against them poses huge challenges [3].

Computer simulations, especially molecular dynamics (MD) simulations have become essential tools to investigate the relationship between conformational and structural properties of proteins and the intermolecular interactions that give rise to aggregation [2,4,5]. In the 21^st^ century, powerful supercomputers have enabled us to simulate more and more complex systems for longer time scales and larger length scales in order to approach experimental conditions.

However, MD simulations generate a large amount of data and extracting information about the relevant molecular processes from them requires adequate post-processing techniques. One of these techniques are Markov state models (MSMs), which have recently gained importance in the fields of computational biochemistry and biophysics as a technique for elucidating the relevant states and processes hidden in the MD data [6,7]. MSMs are network models that encode the system dynamics in a states-and-rates format, i.e., the molecular system can exist in one of many possible states at a particular point in time, which has a fixed probability of transitioning to other states, including itself, within a particular time interval. A basic assumption of MSMs is memorylessness, i.e., the probability of transition from one state to another depends only on the current state and not the history of the system. The suitability of MSMs for extracting essential information from MD data was demonstrated for a large range of biological systems, including protein folding [8], protein-ligand binding [9], or allostery [10]. Our group recently extended the applicability of MSMs to molecular self-assembly by accounting for the degeneracy of aggregated during the aggregation process [11]. The power of this approach for the elucidation of kinetically relevant aggregation pathways has been demonstrated for the self-assembly of the amyloidogenic peptide KFFE [11].

An alternative network model to characterize protein aggregation is provided by transition networks (TNs), which were also developed by the Strodel lab [12]. TNs are based on conformational clustering, instead of kinetic clustering as done in MSMs. In TNs, the aggregation states are defined based on characteristics that are found to be most relevant for describing the aggregation process under study. These so-called descriptors always include the aggregation size and are augmented by, e.g., the number and type of interactions between the proteins in the aggregates, their shape and amount of β-sheet, i.e., quantities relevant to amyloid aggregation. The transformation of the high-dimensional conformational space into this lower-dimensional TN space enables clear views of the structures and pathways of the aggregation process. We successfully applied this approach to the aggregation of the amyloid-β peptide (Aβ_42_) connected to the development of Alzheimer’s disease [13-15], a segment of this peptide, Aβ_16-22_ [12,16], as well as GNNQQNY, a polar peptide sequence from the yeast prion protein Sup35 [12].

In this chapter we provide a guided manual for performing MD simulations of protein aggregation, and analyzing them either with Markov state models or with transition networks.

## 2 Simulation and Analysis Protocols

The basic prerequisite to perform MD simulations of proteins is an MD software engine such as GROMACS [17], AMBER [18], or NAMD [19]. Here, we employ the GROMACS software to illustrate the setup, conductance, and analysis of protein aggregation simulations. There are few more software packages which will be required for the following protocols: 1) protein visualization programs, i.e., PyMOL [20] or VMD [21], 2) Python [22] for general data analysis, 3) Python libraries specifically designed to analyze MD trajectories, i.e., MDAnalysis [23] and MDTraj [24], and 4) a molecule packing optimization software, i.e., PACKMOL [25].

### 2.1 MD Simulations and basic analysis

Most of the following protocols use Aβ_16-22_ as an example, which is treated with capping groups at both ends and thus has the sequence ACE-KLVFFAE-NME. As the Protein Data Bank (PDB) does not include a structure for this peptide, a starting structure for the following simulations can be retrieved from the coordinates of residues 16-21 of a PDB structure of Aβ_42_, as given by the PDB entry 1Z0Q [26]. Using the *Builder* tool in *Protein* mode of PyMOL, the ACE and NME capping groups can be added to the N- and C-terminus, respectively. In this protocol, six copies of Aβ_16-22_ are simulated employing GROMACS 2016.4 as the MD engine, Charmm36m as the protein force field [27], and the TIP3P water model [28]. We use the Charmm36m force field as since it has been shown to be one of the best force fields for modeling Aβ [29] and which also performs the best in our in-house peptide aggregation benchmark [16].

#### 2.1.1 Preparation of the simulation box containing six peptides

1. The first step is to produce a relaxed conformation for the Aβ_16-22_ monomer. This can be achieved with an MD simulation of the monomer following our MD protocol published in Ref. [30]. Alternatively, the MD online tutorial available on our group website can be used: http://www.strodel.info/index_files/lecture/html/tutorial.html. The length of the simulation depends on the size of the peptide under study, for Aβ_16-22_ a simulation length of 1 μs or longer is recommended. The most stable monomer structures can be determined using conformational clustering [31] and six of these structures are used to build the initial system of six Aβ_16-22_ monomers randomly placed in a simulation box. The initial simulation of the monomer is performed to avoid aggregation of artificial peptide structures in the following step, which would require more simulation time for relaxation of such aggregates or, even worse, might lead to artefacts in the simulation data.
2. To randomize positions of the six monomers in a simulation box, we use PACKMOL. The sample script below places six Aβ_16-22_ peptides with at least 1.2 nm (or 12 Å as in the script) distance between them in a simulation box of size ∼10 nm x 10 nm x 10 nm.

~~~
#Six monomers of abeta16-22 peptide
#minimum distance between two monomers
tolerance 12.0
seed −1
#The file type of input and output files is PDB
filetype pdb
#The name of the output file
output abeta16-22_hexamer.pdb
#add TER to after each monomer chain
add_amber_ter
#distance from the edges of box
add_box_sides 1.0
#path to input structure file
#units for distance is measured in Angstrom
#box size is 100 Å
structure abeta16-22.pdb
 number 6
 inside box 0. 0. 0. 100. 100. 100.
end structure
~~~

Alternatively, one can use GROMACS to achieve the same goal:

~~~
gmx insert-molecules -ci abeta16-22.pdb -nmol 6 -box 10 10 10 -o abeta16-22_hexamer.pdb
~~~

As an illustration, the resulting simulation box is shown in Fig. 1.

**Figure 1:**
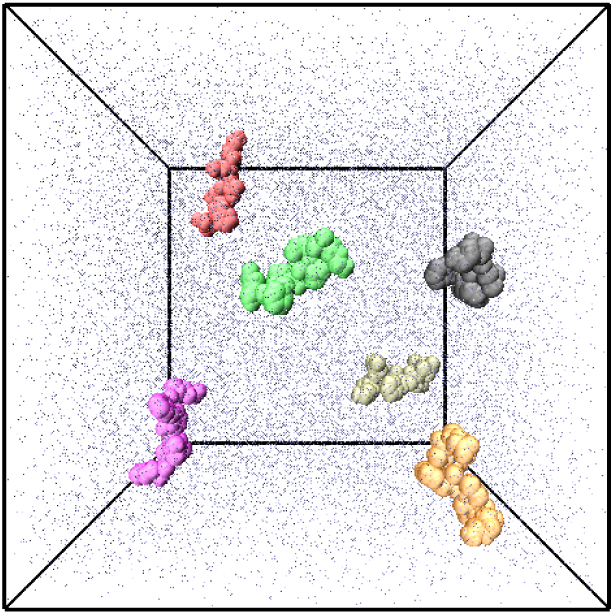
An illustrative example of six Aβ_16-22_ peptides (shown as surfaces in different colors) randomly placed in a cubic box surrounded by solvent molecules (shown as grey dots).

#### 2.1.2 Creation of directories for the different simulation steps

For the execution of the following MD steps, it is advantageous to perform them in separate directories, which avoids accidental replacement of files. To this end, directories for the five major steps are created: topology building, energy minimization, NVT equilibration, NPT equilibration, and MD production run.

~~~
mkdir 1-topol 2-em 3-nvt 4-npt 5-md
~~~

For each step an .*mdp* file is required. The *mdp* file type extension stands for molecular dynamics parameters as these files contain all the key parameters to set up an MD simulation with GROMACS. The five .*mdp* files required are provided in Appendix A at the end of this chapter. Create a directory,

~~~
mkdir mdp
~~~

and copy these .*mdp* files to that directory.

#### 2.1.3 Topology building

In this step the topology file is created. It contains information about molecule types and the number of molecules, which will be simulated. As input, the .*pdb* file from the previous step is taken and, in addition to the topology file, a .*gro* file is produced, which, like a .*pdb* file, also contains the coordinates of the simulated system. The main difference between them is their format. Moreover, a .*gro* file can also hold velocities.

1. Download the Charmm36m force field from http://mackerell.umaryland.edu/download.php?filename=CHARMM_ff_params_files/charmm36-mar2019.ff.tgz and copy it to the *1-topol* directory. Change to that directory:

~~~
cd 1-topol/
~~~
2. Run the GROMACS *pdb2gmx* command to process the input structure file and create the topology file with .*top* extension, topology include files with .*itp* extension, and position restraint files with .*itp* extension.

~~~
gmx pdb2gmx -f ../abeta16-22_hexamer.pdb -o protein.gro -p topol.top -ignh -ter <<EOF
1
1
3
4
3
4
3
4
3
4
3
4
3
4
EOF
<u>Explanation of flags and options:</u>
-f: reads the input structure file *abeta16-22_hexamer.pdb*
-o and -p: writes the output structure file *protein.gro* and system topology file *topol.top*
-ignh: ignores the hydrogen atoms in the input file, which is advisable due to different naming conventions
of hydrogen atoms in input files and force fields. New hydrogen atoms will be added by
GROMACS using the H-atom names of the selected force field.
-ter: to interactively assign charge states for N- and C-terminal ends
Option 1: choosing protein force field (charmm36-mar2019)
Option 1: choosing water force field (TIP3P)
Option 3: choosing N-terminus (None, as we use ACE capping)
Option 4: choosing C-terminus (None, as we use NME capping)
Options 3 and 4 are repeated for each peptide in the system, in this example six.
~~~ After a successful execution of the GROMACS *pdb2gmx* command, the directory will also contain six topology-include files and six position restraint files, one for each peptide: *to-pol_Protein_chain_$chain.itp* and *posre_Protein_chain_$chain.itp* with $*chain* = *A, B, C, D, E*, or *F*. The latter files contain position restraint entries for all non-hydrogen atoms of the peptides, which are needed during the equilibration MD steps.
3. Next, a simulation box is created. Note that the box defined above was only needed for placing the peptides. In this step, the MD simulation box is set up. As before, a cubic box of 10 x 10 x 10 nm^3^ is chosen:

~~~
gmx editconf -f protein.gro -o box.gro -bt cubic -box 10 10 10
~~~
4. Now the simulation box is being filled with water, solvating the peptides. For this an existing patch of 216 water molecules is used (*spc216.gro*) and repeated in *x, y*, and *z* direction until the box is completely filled, thereby making sure that water and peptides do not overlap.

~~~
gmx solvate -cp box.gro -cs spc216.gro -o protein-solvated.gro -p topol.top
~~~ The addition of solvent molecules is reflected in the *topol.top* file, which now includes 32,323 water molecules in addition to the six peptides.
5. For performing an MD simulation with periodic boundary conditions (PBCs), we need to neutralize the charge of the system by adding positive (Na^+^) or negative (Cl^-^) ions as needed in order to avoid artefacts during the calculation of the electrostatic interactions. Also, we can assign a specific ion concentration to the system, which is often ∼150 mM to mimic physiological conditions:

~~~
gmx grompp -f ../mdp/ions.mdp -c protein-solvated.gro -p topol.top -o protein-ions.tpr
echo 13 | gmx genion -s protein-ions.tpr -o protein-ions.gro -p neutral -conc 0.15
<u>Explanation of flags and options:</u>
-neutral: neutralizes the system to charge zero by adding Na^+^ or Cl^-^ ions
-conc: changes the concentration to 150 mM NaCl
echo 13: replaces the solvent molecules with ions where option 13 stands for SOL (solvent molecules).
~~~

In the first command, the *grompp* module, also known as GROMACS pre-processor, reads the coordinate file, the system topology file, the *ions.mdp* file and processes them into a GROMACS binary format generating a .*tpr* file, where the file extension stands for portable run input file. This file contains the starting structure for the simulation, the topology information of the individual peptides and water molecules, as well as all the simulation parameters. The second command reads the binary input file *protein-ions.tpr* and neutralizes the net charge – if the overall charge of peptides should not be zero, which is not the case in our current example – and increases the salt concentration of the system to the specified value by replacing solvent molecules. This results in 90 Na^+^ and 90 Cl^-^ ions being added in our example; the *topol.top* file is accordingly updated.

Now everything is prepared to start with the equilibration of the system, consisting of an initial energy minimization and two short MD simulations.

#### 2.1.4 Energy minimization

1. For the energy minimization (EM) step, change to the directory *2-em*:

~~~
cd ../2-em
~~~
2. As indicated in the *em.gro* file, we employ the steepest descent algorithm to minimize the system until a maximum force of 500 kJ/mol/nm or 10,000 minimization steps are reached. The EM is performed to ensure that the MD can be started, for which relatively small forces are needed.

~~~
gmx grompp -f ../mdp/em.mdp -c ../1-topol/protein-ions.gro -p ../1-topol/topol.top -o protein-em.tpr
~~~ Once again, *grompp* is used to combine the structure, topology, and simulation parameters into a binary input file, *protein-em.tpr*, which is then passed to the GROMACS *mdrun* command.
3. The *mdrun* command is invoked by

~~~
gmx mdrun -v -deffnm protein-em
<u>Explanation of flags:</u>
-v: verbose, prints the progress of EM step to the screen after every step
-deffnm: defines the input and output filenames
~~~ On successful execution of the *mdrun* command, following files are generated:

~~~
protein-em.gro: final energy.minimized structure file
protein-em.edr: energy file in binary format
protein-em.trr: trajectory file including all the coordinates, velocities, forces, and energies in binary format
protein-em.log: text log file of EM steps in ASCII format
~~~

#### 2.1.5 NVT Equilibration

Following the EM step, two equilibration (EQ) steps are performed. The first EQ step is conducted under isothermal and isochoric conditions, which are called *NVT* ensemble as *N* (the number of particles), *V* (the volume), and *T* (the temperature) are held constant. Moreover, at this step only the solvent molecules and ions get equilibrated around the peptides, bringing them to the desired temperature (300 K in our case), while the positions of the peptide atoms are restrained. To this end, the .*itp* files containing the position restraints, which were generated during topology building, are used. Similar to the EM step, in the *NVT* and the following MD steps *grompp* and *mdrun* are called:

~~~
cd ../3-nvt
gmx grompp -f ../mdp/nvt.mdp -c ../1-topol/protein-em.gro -p ../1-topol/topol.top -o protein-nvt.tpr
gmx mdrun -v -deffnm protein-nvt
~~~

Typically, the *NVT* EQ step is a 100-ps long MD simulation, which suffices to equilibrate the water around the proteins or peptides at the desired temperature. Here is an explanation of important parameters set in the *nvt.mdp* file:

~~~
gen_seed = −1: takes as random number seed the process ID.
gen_vel = yes: generates random initial velocities. For this, the random number seed is used.
tcoupl = V-rescale: defines the thermostat.
pcoupl = no: pressure coupling is not applied.
~~~

If *grompp* assigns a random number seed based on its process ID, every time one re-runs *grompp* it will assign a different seed, because the respective process ID for that grompp execution is also different. This guarantees that the seeds will always be random so that each time the simulation is repeated, different random numbers are generated, leading to a different initial velocity distribution. This is important when a simulation is repeated several times to collect statistics on a system. The temperature coupling is achieved using a velocity rescaling thermostat [32], which is an improved Berendsen weak coupling method. Upon successful execution of the *mdrun* command, files with the same file type extensions as generated in the EM step are produced.

#### 2.1.6 NPT Equilibration

In the second EQ phase, the pressure and thus density of the system are adjusted using the iso-thermal-isobaric ensemble, also called or *NpT* ensemble as *N, p* (the pressure), and *T* are kept constant. The 200-ps *NpT* EQ simulation is executed with the help of the *grompp* and *mdrun* commands:

~~~
cd ../4-npt
gmx grompp -f ../mdp/npt.mdp -c ../1-topol/protein-nvt.gro -p ../1-topol/topol.top -o protein-npt.tpr
gmx mdrun -v -deffnm protein-npt
~~~

One change in the *npt.mdp* file compared to the *NVT* EQ run is the addition of the pressure coupling section, using the Parrinello-Rahman barostat [33]. The other notable changes are:

~~~
continuation = yes: continuation of the simulation from the *NVT* EQ step.
gen_vel = no: velocities will be read from the trajectory files generated from the NVT equilibration step
and not newly initiated.
~~~

After successful execution of the EQ phases the temperature and pressure of the whole system are adjusted, so that we can proceed to perform the production run.

#### 2.1.7 MD Production Run

In the production run, the position restraints on the proteins are removed; however, all bond lengths will be constrained to their equilibrium values using the LINCS method [34] which allows to use a time step of 2 fs for the integration of the equations of motions. Otherwise, the MD production run is similar to the *NpT* EQ step. To sufficiently sample the conformational space, we need to run production runs in the order of microseconds.

To perform the 1-μs MD production run in this example, change into the corresponding directory, call the GROMACS pre-processor, *grompp*, and then the *mdrun* command:

~~~
cd ../5-md
gmx grompp -f ../mdp/md.mdp -c ../1-topol/protein-npt.gro -p ../1-topol/topol.top -o protein-md.tpr
gmx mdrun -v -deffnm protein-md
~~~

After the MD production run, another file type compared to the previous steps will be generated, an .*xtc* file, which results from a directive in the *md.mdp* file concerning the output parameters. An .*xtc* file is a portable format that saves trajectories using a reduced precision.

#### 2.1.8 MD analysis: Oligomer size and contact maps

After successful completion of the MD production run, analysis of the MD trajectory can commence. The *protein_md.xtc* file harbors all the coordinates of the system sampled by the production simulation that are used for the analysis. The various GROMACS analysis tools can read the binary format of the .*xtc* files. However, in the following we will employ the Python-based MDAnalysis and MDTraj like tools, which can also handle .*xtc* files, in connection with our in-house Python scripts. For the current example of the system with six Aβ_16-22_ peptides we will limit ourselves to two quantities to be analysed: the oligomer size and inter-peptide contacts in the aggregates.

1. Before we embark on starting the analysis, create a new directory for the analysis:

~~~
cd ../
mkdir analysis
cd analysis/
~~~ Copy the trajectory file *protein_md.xtc* and the run input file *protein_md.tpr* from the production MD directory to the analysis directory.
2. For the analysis, we only need the coordinates of the proteins, but not those of the solvent and ions. The extraction and re-saving is performed using the GROMACS *trjconv* command:

~~~
gmx trjconv -s protein_md.tpr -f protein_md.xtc -o protein_only.xtc
<u>On the command prompt:</u> select
option “1” for centering “protein”,
option “1” for output “protein”
The output .*xtc* file will include only coordinates of the peptide chain.
~~~
3. We also need to extract a reference structure file either in .*gro* or .*pdb* format. In the current example, we create a .*pdb* file.

~~~
gmx trjconv -s protein_md.tpr -f protein_md.xtc -o protein_only.pdb -dump 0
<u>On the command prompt:</u> select
option “1” for output “protein”
<u>Explanation:</u>
-dump 0: dumps the first frame of the trajectory file.
~~~
4. A particular problem of protein aggregation simulations that needs to be addressed prior analysis are the PBCs during the MD simulation that can cause the proteins appearing to be broken. Many of the analysis scripts cannot handle such broken proteins, which would lead to artefacts in the analysis. To this end, we have to revert the effects of the PBCs prior the analysis, for which we use VMD. Read in the newly created *protein_only.pdb* and *protein_only.xtc* files in VMD and visualize the trajectory. Then open the *Tk Console* on the *Extensions* tab and visualize the PBC box around the protein system by entering the command:

~~~
pbc box
~~~ When playing the trajectory movie, one can see some frames having broken molecules, which can be reassembled by entering following commands in the Tk Console:

~~~
[atomselect top all] set chain 0
pbc join fragment -all
~~~ Afterwards, save *all* coordinates to a new name in the analysis directory as *protein_nopbc.trr.* This trajectory file will be used in the following analysis.
5. The Python scripts to calculate the oligomerization state (monomer, dimer, trimer etc.) and the inter-residue contact frequencies between peptides or proteins are available in Appendix B.

The script is automated and requires *protein_only.pdb* and *protein_nopbc.trr* as input files and generates following output:

~~~
#Input files
protein_only.pdb
protein_nopbc.trr
#Important output files
Oligomer_groups.dat: groups the interacting protein chains.
Oligomer_states.dat: counts the number of chains in each oligomer group.
Oligo-highest-size.dat: finds the maximum oligomer size formed per MD frame.
Oligo-block-average.dat: creates 25-frame moving averages of the maximum oligomer size.
Contact_map.dat: saves the frequency of contacts between residues from different proteins.
~~~

The results obtained from this analysis applied to our example are shown in Fig. 2. Panel A reports on the oligomerization state of system, where two proteins were considered to be in contact with each other if the minimum distance with respect to any two atoms from either protein was below 0.4 nm. Only the maximum oligomer size in the system at a given time is reported. For example, if at one point the system consists of a dimer and a tetramer, then the tetramer as the larger oligomer is of interest. The plot in Fig. 2A shows that the six peptides reached the hexamer state within ∼300 ns, but a few dissociation and reassociation events are observed at later times, especially between 500 and 750 ns. The residue-residue contacts between the peptides composing the oligomers are then counted and reported as probability map in Fig. 2B. It shows the peptides prefer to assemble in an anti-parallel orientation, which are stabilized by electrostatic interactions between the oppositely charged N-terminal K16 and C-terminal E22 residue. In addition, a few strong hydrophobic contacts are formed, especially L_*i*_17-F_*j*_19, V_*i*_18-V_*j*_18, and F_*i*_19-L_*j*_17 where *i* and *j* refer to two different peptides in an oligomer.

**Figure 2:**
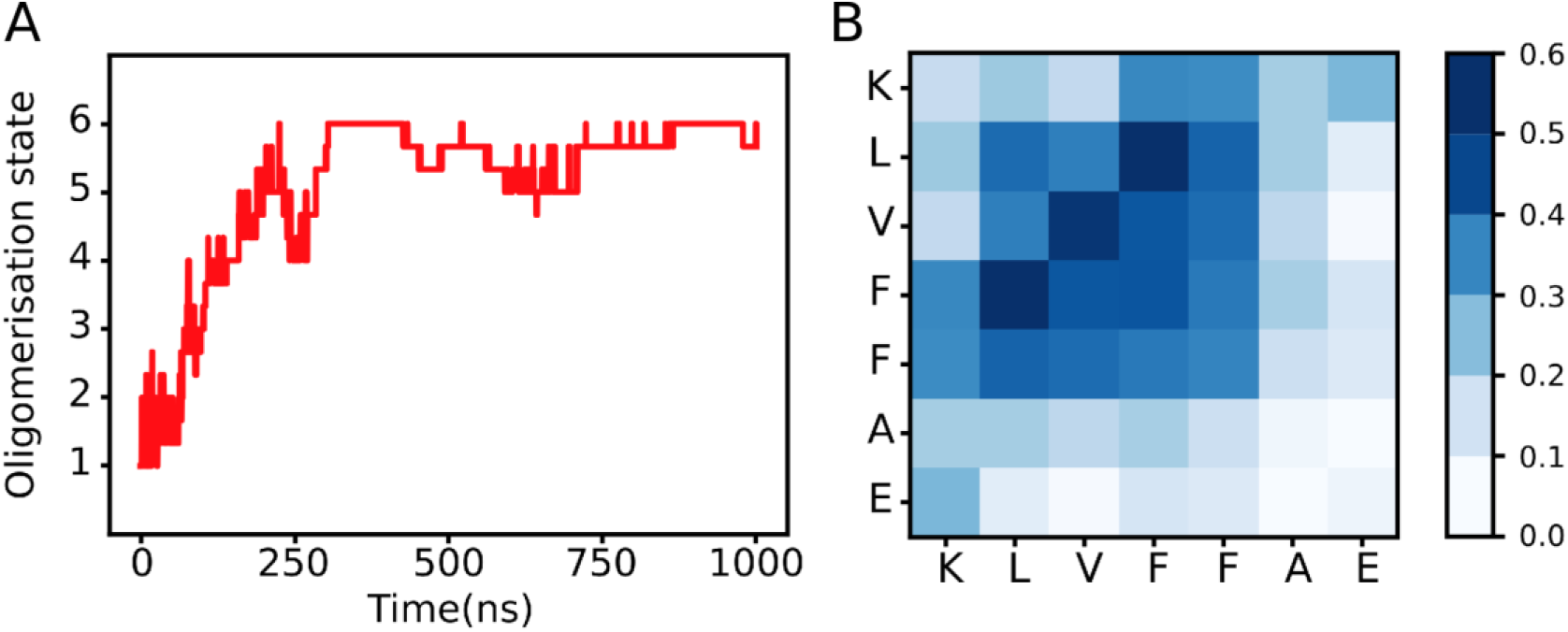
Analysis of the 1-μs MD simulation following the aggregation of six Aβ_16-22_ peptides. **A**) The oligomerization state of the system over time is shown. **B**) The inter-residue contact map with probabilities according to the color scale on the right is shown. The contacts between pep- tides composing the oligomers sampled during the simulation are reported.

Our in-house Python scripts are available at https://github.com/strodel-group/Oligomerization-State_and_Contact-Map.

### 2.2 Markov state models for the analysis of protein aggregation

Markov state modelling, is a mathematical framework for time-series data analysis, which can be used for understanding the underlying kinetics hidden in high-dimensional MD simulation data. Specifically, it enables the construction of an easy to interpret states-and-rates picture of the system consisting of a series of states and transition rates between them. This tutorial is a short introduction to the construction of Markov state models from MD trajectory data with the help of the PyEMMA library [35] in Python.

The first step towards building a Markov state model (MSM) using PyEMMA is to choose a suitable distance metric for defining the feature space of the system, followed by reducing the dimension of this space using a suitable dimensionality reduction technique. Here, the method of choice is usually time-lagged independent component analysis (TICA) [36]. In simulations of molecular self-assembly as in the current example one has to accounting for the degeneracy of oligomers during the aggregation process, which results from numbering the identical molecules during the simulation. Our lab solved this problem by sorting the permutable distances of the feature space, which we implemented into TICA and is available as TICAgg (TICA for aggregating systems, https://github.com/strodel-group/TICAgg) [11]. Next, some clustering method is applied decomposing the reduced conformation space into a set of disjoint states, which are then used to transform the trajectory into a sequence of transitions between these states. An MSM can be built from this discrete trajectory by counting the transitions between the states at a specified lag time, constructing a matrix of the transition counts, and normalizing it by the total number of transitions emanating from each state to obtain the transition probability matrix. The Markovian character of the model can be verified with the Chapman-Kolmogorov test. However, the model is often too granular to provide a simple, intuitive picture of the system dynamics. That is achieved by coarse-graining the model into a hidden Markov model (HMM) with a few metastable states, using robust Perron cluster analysis (PCCA+) [37].

The quality and practical usefulness of an MSM mainly depends on the state-space discretization, which includes feature selection, dimensionality reduction, clustering, and the selection of the lag time of the model. Hence, for obtaining an MSM that is both descriptive and predictive, an appropriate way for adjusting the hyperparameters in each step of the PyEMMA workflow is necessary. In the following we provide guidelines for these selections using the aggregation of Aβ_16-22_ into a dimer as example. For this example, we simulated the system for 10 μs in total. As Markov state modeling can be combined with adaptive sampling, it is not necessary to simulate one long trajectory. Instead, several short trajectories can be used for the MSM analysis, which are ideally started from different and initially rarely sampled states; hence the name “adaptive sampling”.

#### 2.2.1 Feature Selection

1. First, a featurizer object, which will hold information about the system’s topology, has to be generated by loading the topology file for every feature, e.g. backbone torsion angles:

~~~
torsionsFeat = pyemma.coordinates.featurizer(topologyFile)
~~~
2. Next, the featurizer object is initialized with its feature:

~~~
torsionsFeat.add_backbone_torsions()
~~~ When studying protein aggregation, the system is often composed of several identical chains, which can lead to degenerate aggregates. However, in simulations they are usually not recognized as being identical due to molecular indexing. To resolve this issue, we sort the permutable distances as implemented in TICAgg [11] and define intermolecular distances and intramolecular distances as further features:

~~~
interDistFeat = pyemma.coordinates.featurizer(topologyFile)
interDistFeat.add_custom_func(dist_intermol, dim_permute_sortinter)
intraDistFeat = pyemma.coordinates.featurizer(topologyFile)
intraDistFeat.add_custom_func(dist_intramol, dim_permute_sortintra)
~~~ Note: When working with monomeric systems, intermolecular distances should not be used as feature and the usual TICA procedure should be employed, while the workflow of TICAgg should be followed for aggregation studies (https://github.com/strodel-group/TICAgg).
3. Load the MD simulation data from the trajectory files (how many there are depends on the user, see comment above) by using the featurizer objects:

~~~
torsionsData = pyemma.coordinates.load(trajectoryFiles, features=torsionsFeat)
interDistData = pyemma.coordinates.load(trajectoryFiles, features=interDistFeat)
intraDistData = pyemma.coordinates.load(trajectoryFiles, features=intraDistFeat)
~~~
4. Next, find the best suited feature for capturing the slow dynamical processes. Here, the VAMP-2 score [38] is an established measure for ranking them. The aim is to extract the feature which preserves the most kinetic variance corresponding to the highest VAMP-2 score:

~~~
vamp2 = pyemma.coordinates.vamp(data=torsionsData[:-1], lag=100,
dim=10).core(test_data=torsionsData[-1], score_method = ‘VAMP2’)
~~~

In the case shown above, the entire dataset of trajectories is used for learning except for the last trajectory (visible from the indexing “[:-1]”) which is used as validation set for calculating the VAMP-2 score. It is recommended to compute the score for a bunch of learning and validation sets with varying compositions. Also, the VAMP-2 score should be determined for several lag times *lag* (in units of the trajectory time step). The number of dimensions *dim* is fixed in order to guarantee the comparability of different features.

The VAMP-2 scores of the different features considered for the current example are presented for several lag times in Fig. 3.

**Figure 3:**
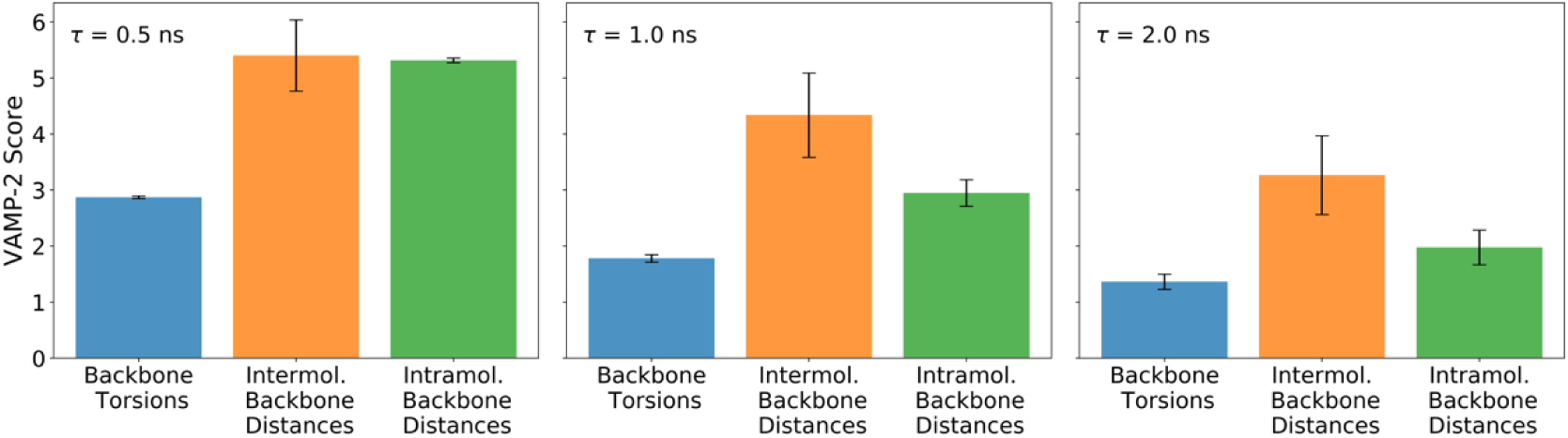
The VAMP-2 obtained for different features and lag times derived from a 10-μs MD simulation of two Aβ_16-22_ peptides.

Obviously, the intermolecular backbone atom distances are superior for describing the system’s dynamics at all lag times, as the VAMP-2 score of this feature is always the highest. Therefore, intermolecular backbone atom distances are used as feature for the further procedure.

#### 2.2.2 Dimension reduction and discretization

The feature space is usually high dimensional. Discretizing in such high dimensional spaces is inefficient and would produce low-quality results. Therefore, it is convenient to first reduce the dimension of the selected feature space before discretizing it. Here, TICAgg is used as technique for dimensionality reduction and k-means clustering for partitioning the reduced space.

1. Since TICA works by attempting to maximize the autocorrelation of the given coordinates, the lag time should be in the order of magnitude of the desired processes time scales. Here, a lag time of 5.0 ns (*lag* = 250, which corresponds to the number of frames considering that every 20 ps a frame was saved in the trajectory) is selected. The number of dimensions to keep should be as small as possible, but high enough to capture the important events. Here, we keep 10 dimensions (*dim* = 10).

~~~
tica = pyemma.coordinates.tica(interDistData, lag=250, dim=10)
ticaOutput = tica.get_output()
~~~
2. The sample density in this reduced space can be revealed by plotting the data along the main TICA components (called ICs). For visualization purpose, the *ticaOutput* of the trajectories is concatenated.

~~~
ticaOutputConcatenated = np.concatenate(ticaOutput)
IC1 = ticaOutputConcatenated.T[0]
IC2 = ticaOutputConcatenated.T[1]
pyemma.plots.plot_density(IC1, IC2, logscale=True)
~~~ The resulting plot for the Aβ_16-22_ dimer is shown in Fig. 4. **Figure 4:**
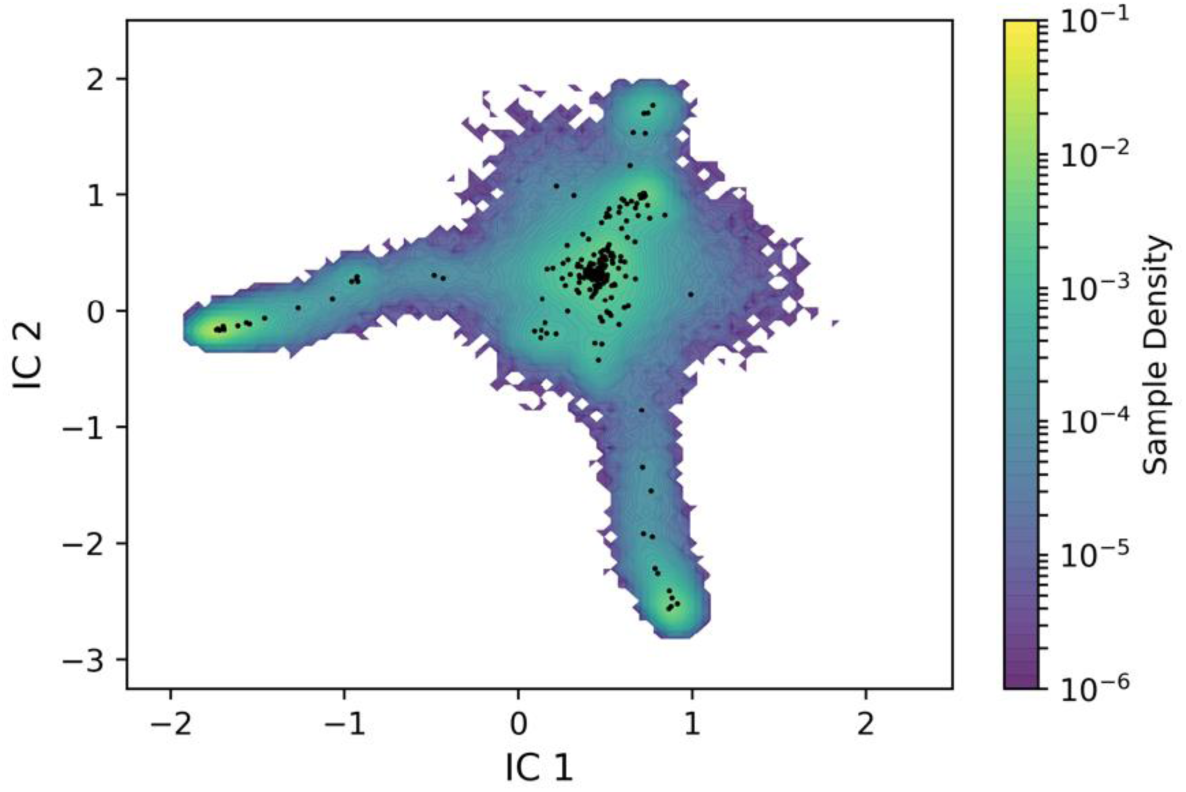
Sample density, according to the color scale on the right, along the first two ICs obtained from TICA applied to the 10-μs MD data of two Aβ16-22 peptides. The black dots represent the location of the 200 cluster centres in the IC1-IC2 resulting from k-means clustering.
3. The next step is to discretize the trajectories in TICA space using k-means clustering for defining microstates. Here, a balance between low computational effort (minimize the number of cluster centres) and high preservation of dynamic information content (maximize the VAMP-2 score) is wanted. This optimization problem is solved by analysing the VAMP-2 score in dependency of the number of cluster centres *nClusterCentres*. For the system under investigation, the number of cluster centres is set to 200 and the result can be seen in Fig. 4.

~~~
nClusterCentres = 200
cluster = pyemma.coordinates.cluster_kmeans(ticaOutput, k=nClusterCentres, max_iter=100)
~~~
4. Finally, the discrete trajectories are generated using the cluster object.

~~~
discretizedTrajectories = cluster.dtrajs
~~~

#### 2.2.3 Construction of the Markov State Model

1. A lag time for the construction of the Markov state model has to be selected. As the best lag time is initially not known, one generates MSMs for different lag times *lagTimes* and the dependency of their *N* implied time scales (ITs) on the lag time is studied.

~~~
lagTimes = [lagTime for lagTime in range(2, 200)]
lagTimes.extend([300, 500, 1000])
N = 20
impliedTimeScales = pyemma.msm.its(discretizedTrajectories, lags=lagTimes, nits=20)
~~~
2. The lag time dependency of the implied time scales can be visualized (see Fig. 5) using a PyEMMA plotting tool:

~~~
pyemma.plots.plot_implied_timescales(impliedTimeScales)
~~~

**Figure 5:**
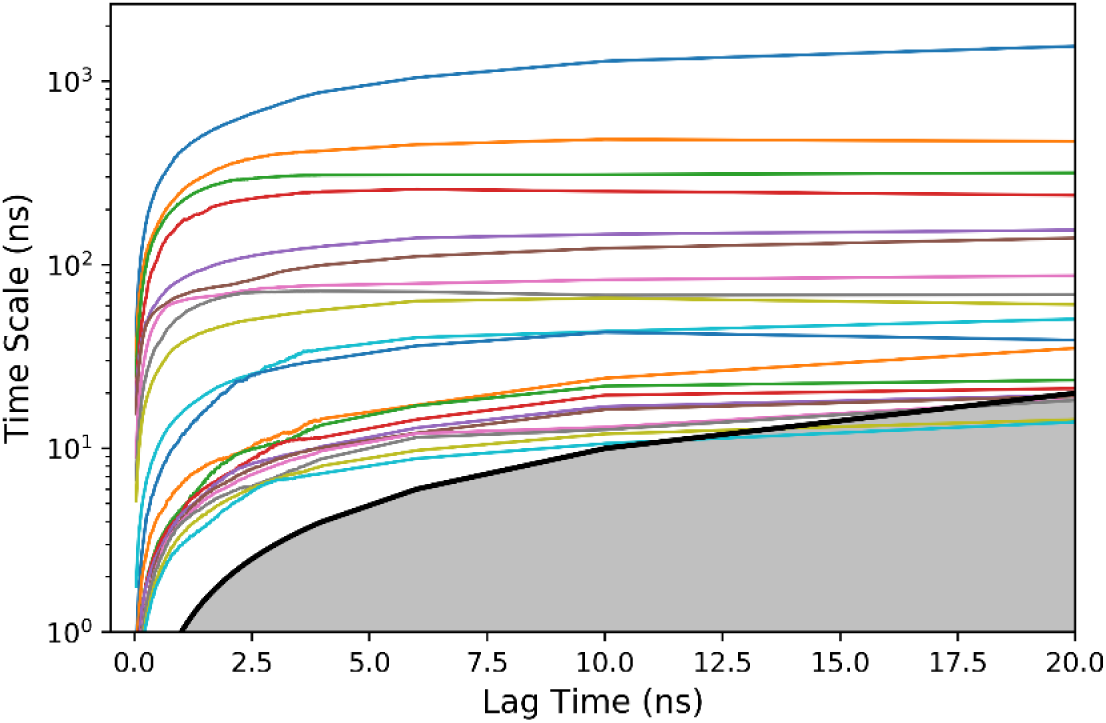
Implied time scales of the 20 slowest processes for different Markov state models at different lag times (*x*-axis) of the Aβ_16-22_ dimer system. The black line separates the area where the dynamics of the processes is resolvable (white) from the non-resolvable area (grey).
3. To ensure that the ITs are lag-time independent, which is necessary for the Markovianity of the model, the focus is directed on areas in Fig. 5 where the graphs are converged. Furthermore, it is important that the lag time to choose for the MSM does not exceed the time scale of the main processes, as otherwise the processes cannot be resolved anymore. This holds true for the white area above the black line in Fig. 5. To keep as much as detail for the dynamics as possible, the lag time should be chosen from a region where the ITs have just converged. Taking these considerations into account, a lag time of 6.5 ns (*lagTime* = 325) is chosen in our example.

~~~
lagTime = 325
msm = pyemma.msm.estimate_markov_model(discretizedTrajectories, lagTime)
~~~
4. To verify that the model is Markovian, a Chapman-Kolmogorov test is performed. Since this validation routine requires macrostates rather than microstates as input, the number of aspired coarse-grained states *nStates* has to be specified first. A reasonable criterion for this choice is the separation of two consecutive implied time scales, which should be high to enable a clear distinction between slow and fast processes. For the Aβ_16-22_ dimer especially high separation values are present between the fourth and fifth time scale, leading to five macrostates to coarse-grain into.

~~~
nStates = 5
chapmanKolmogorov = msm.cktest(nStates)
pyemma.plots.plot_cktest(chapmanKolmogorov)
~~~

Fig. 6 shows the Chapman-Kolmogorov test of the constructed MSM. It can be seen that all predicted transition probabilities (black continuous lines) agree very well with the estimated ones (blue dashed lines), which confirms the Markovianity of the model.

**Figure 6:**
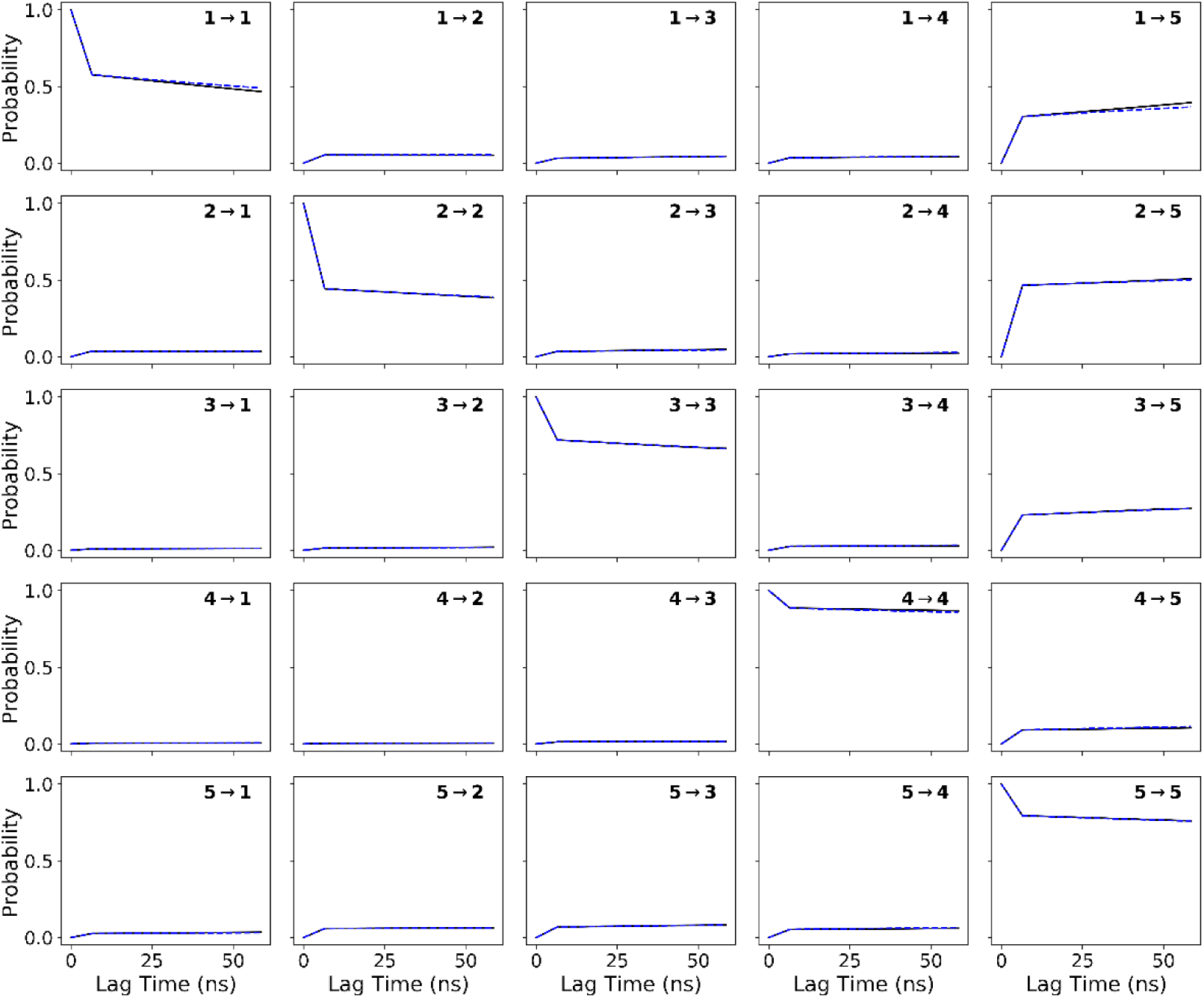
Chapman-Kolmogorov test for the constructed Markov state model (*lag* = 325, *nStates* = 5) of the Aβ_16-22_ dimer system. Estimations (blue) and predictions (black) are shown.

#### 2.2.4 Hidden Markov Model

MSM models generally have a few hundreds to thousands number of microstates (resulting after the k-means clustering step) and are as such too granular to provide a human readable network and thus an easy to comprehend picture of the system. To this end, the MSM needs to be coarse-grained into a given number of macrostates using the Perron Cluster Analysis method (PCCA+) [37]. The resulting model is known as hidden Markov model.

1. Specify the number of macrostates *nStates* to construct the HMM:

~~~
hmm = msm.coarse_grain(nStates)
~~~
2. Finally, the transition matrix can be visualized as a network:

~~~
pyemma.plots.plot_markov_model(hmm)
~~~

The HMM resulting from our example is shown in Fig. 7. It is overlaid onto the sample density from Fig. 4 and complemented by a representative structure for each of the five macrostates. Macrostate 5 is predominantly composed of monomeric Aβ_16-22_ structures, while the four other states correspond to dimeric structures. They are all antiparallel β-sheets, which differ in their registry and thus inter-residue contacts and amount of β-sheet. State 4 is the in-register antiparallel β-sheet with contacts L_1_17-A_2_21 and F_1_19-F_2_19. State 2 is similar to state 4, yet the β-sheet is shorter. States 1 and 3 are out-of-register antiparallel β-sheets which differ by their inter-peptide contacts: L_1_17-F_2_19 and F_1_19-L_2_17 in state 1 and K_1_16-F_2_20, V_1_17-V_2_17, and F_1_20-K_2_16 in state Here, the indices indicate the two peptide chains. Another interesting observation from this MSM is that none of the dimers directly interconverts to one of the other dimers; instead the dimers first dissociate into two monomers before reassociating. This observation agrees to our finding for the short amyloidogenic sequence KFFE and implies that aggregation into β-sheet fibrils most likely proceeds in an orderly manner from the very beginning and not via hydrophobic collapse followed by internal reordering of the aggregates [11]. To our knowledge, such a clear picture about the aggregation pathways can currently only be obtained from MD generated MSMs.

**Figure 7:**
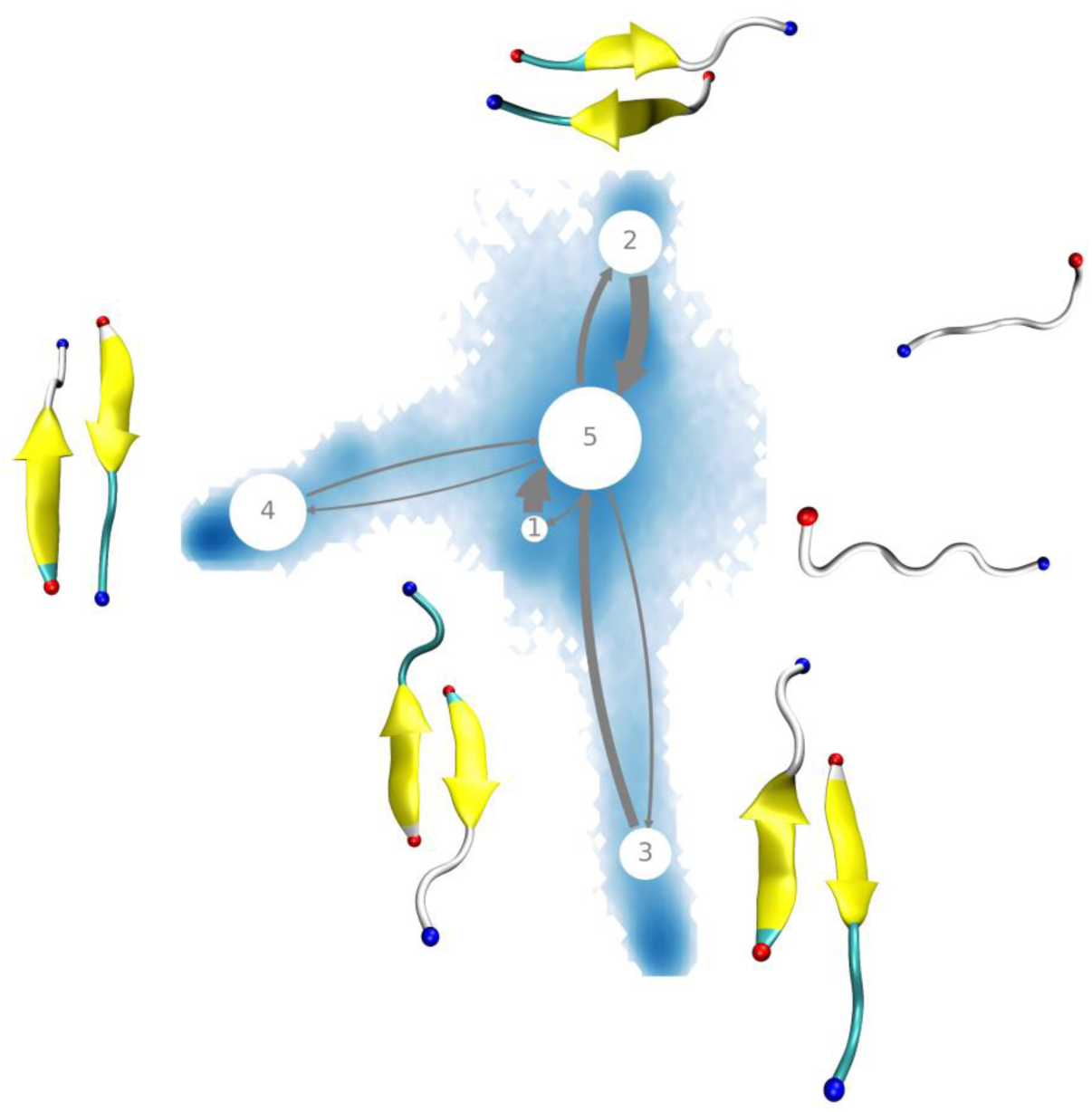
Coarse-grained MSM, also called hidden Markov model, for the Aβ_16-22_ dimer system overlaid on the sample density in the TICA IC1-IC2 space. Representative structure for all five macrostates are shown as cartoon. The N- and C-termini are indicated by blue and red spheres, respectively, while β-sheets are colored in yellow.

### 2.3 Transition networks for the analysis of protein aggregation

Transition networks (TN) are a great analysis tools for studying the assembly of peptides or proteins into oligomers based on MD simulations [12-16]. Once a trajectory is obtained, a transition matrix can be derived that contains information regarding the aggregation states encountered during the simulation and the transitions between different states. The definition of aggregation states depends on the system under study and on the questions to be answered. We usually define the aggregation states as a collection of structural features of a particular monomer or oligomer that is most suited to describe the conformational changes observed during the assembly process. Thus, it can contain: the oligomer size (i.e., the number of peptides in a given assembly), the average number of hydrogen bonds between the peptides that form the oligomer, the average number of amino acids that are in β-strand conformation, the average number of salt bridges or hydrophobic contacts, or the compactness of an oligomer (defined by the ratio of the largest and the smallest moments of inertia).

Here we study the assembly process of two peptides of 25 amino acids each into a dimer using transition networks. The peptide is subrepeat R1 of the functional amyloid CsgA, which was simulated with N-terminal ACE and C-terminal NME capping for 2 μs. CsgA is a functional amyloid secreted by *Escherichia coli* as a soluble protein and aggregates on the plasma membrane upon nucleation by CsgB, aiding in biofilm formation [39]. However, some of the CsgA subrepeats including R1 have been shown to aggregate spontaneously [40]. In this example, dimer formation is analyzed with a TN, but the method can also be applied to higher order oligomers as demonstrated in several of our TN studies [12,13,15,16].

Before starting the TN analysis, a few preparatory steps need to be taken. The analysis script is written in the Tcl scripting language and takes advantage of some useful functions implemented in the VMD software, which we already used in section 2.1 for visualizing the trajectory of Aβ_16-22_ hexamer formation. We assume that an MD trajectory has already been generated. One should make sure that in the trajectory the peptides are complete and not split across the simulation box as a result of PBCs used during the MD simulation. In section 2.1.8 it is explained how reassembled molecules can be achieved. In the following, the trajectory used for analysis is called *md_trajectory.xtc* and the initial atom positions needed by VMD to interpret the .*xtc* file are saved as *md_protein.pdb*.

#### 2.3.1 Running the transition network analysis Tcl script

The analysis will be done by using the Tcl script *TNA.tcl*, which takes care of most of the calculations and is listed in Appendix C. To perform the analysis, one calls the *TNA.tcl* script from VMD with the following command:

~~~
vmd -dispdev text -e TNA.tcl -args md_protein.pdb md_trajectory.xtc 25 2
~~~

VMD is launched in text mode and the arguments needed are the topology or .*pdb* and trajectory files, the number of amino acids in each peptide (25), and the number of peptides in the system (2).

The script first handles the input files, loading the trajectory, renumbering the peptide chains, and extracting the number of frames. Then, within a loop cycling through all frames of the trajectory, the script iterates over all peptides, identifying if they are within the cutoff distance of another monomer or oligomer. Whether the current oligomers decay to monomers or smaller oligomers or form new oligomers is carefully investigated. Transitions between monomers or oligomers from each two consecutive frames are bookkept. Once the transitions at oligomeric level (i.e., monomer, dimer etc.) are uniquely identified, the script proceeds to identify aggregation states for each oligomer by calculating various characteristics of the assemblies. In the current case these characteristics are the number of residues in β-strand conformation (based on the dihedral angles) and the compactness of each oligomer. These specific calculations are included as Tcl procedures at the beginning of the script. In the end, the transition state contains the oligomer size, the average number of amino acids in β-strands, and the compactness. Note that when the procedure returns a value, the main script averages it over the number of peptides in the system and rounds it to the next integer in order to have as few discrete states as possible. A state is then notated by joining the order parameters with a vertical bar as separator, e.g. 1|4|6 which refers to a monomeric state with 4 residues in a β-strand conformation and medium compactness as the last digit can run from 0 for a stick and 10 for a sphere.

Eventually, all recorded transitions between aggregation states are appended to a list and a transition matrix is generated along with the attributes specific to each state. The output files generated contain the state attributes and the transition matrix as described in the following.

#### 2.3.2 The state attributes

The state attributes are saved in the file *State-Attributes.csv*, which lists the identified states and numbers them corresponding to their ID, followed by their oligomer size, residues in β-strand and compactness, and finally gives their population, i.e., the number of their occurrence in the simulation. In the current simulation only 8 states were identified:

~~~
id state oligomer beta-sheet compactness population
1 1|0|2 1 0 2 5
2 1|2|6 1 2 6 1
3 1|2|7 1 2 7 1
4 1|4|6 1 4 6 2
5 1|4|7 1 4 7 1
6 2|1|2 2 1 2 1
7 2|2|2 2 2 2 6
8 2|2|3 2 2 3 1
~~~

#### 2.3.3 The transition matrix

The transitions between the states are saved in the file *Transition-matrix.dat*. In the current example, only eight states were adopted during the trajectory, giving rise to an 8 x 8 matrix (see below). The line numbers (shown in light grey on the left) refer the state ID from where the transition in question occurs, and the column numbers (shown in light grey on the top) are the states into which it occurs. Summing over all entries gives the total number of transitions, which is 18 here. The matrix is not necessarily symmetric because the transition probability from a state *A* to a state *B* can be and in most cases is different from each other. Shown below is the matrix corresponding to this example:

~~~
 1 2 3 4 5 6 7 8
1 3 0 0 0 0 0 2 0
2 0 0 0 0 0 0 1 0
3 0 1 0 0 0 0 0 0
4 0 0 0 0 1 0 1 0
5 0 0 1 0 0 0 0 0
6 0 0 0 0 0 0 1 0
7 1 0 0 1 0 1 1 1
8 0 0 0 0 0 0 0 0
~~~

#### 2.3.4 Visualizing the transition network

The transition matrix can be visualized with the software Gephi (https://gephi.org/). One way to import the matrix into Gephi is to convert the file to a .*csv* format where the values are separated by commas and all rows and columns are numbered starting from 1 to the maximum number of states. The conversion of the transition matrix can be accomplished with the included Tcl script *convert2csv.tcl* and applying the command:

~~~
vmd -dispdev text -e convert2csv.tcl -args Transition-matrix.dat Transition-matrix.csv
~~~

Now the file *Transition-matrix.csv* can be imported into Gephi. The data will be recognized as a matrix, with the first row and first column containing the indices of the transition states that represent the nodes of the network. In the final window of the import section one should select the *Graph Type* as *Directed*, the *Edges merge strategy* as *Don’t merge*, and tick the options *Create missing nodes* and *Self loops*.

The next step is to import the attributes for the network nodes. In the *Data Laboratory* with the *Import Spreadsheet* option one can import the attributes file *State-Attributes.csv*. This data will be recognized as a *Nodes table*. In the last window of the import section the option *Append to existing workspace* should be marked. Note that the indices from the attributes file should correspond to the indices of the imported matrix. Finally, if the size of the nodes is set proportional to the state population, the color of the nodes chosen to correspond to the number of residues in β-strand, the *ForceAtlas 2* layout with non-overlapping nodes and *LinLog* mode selected, one should obtain a figure similar to Fig. 8.

**Figure 8:**
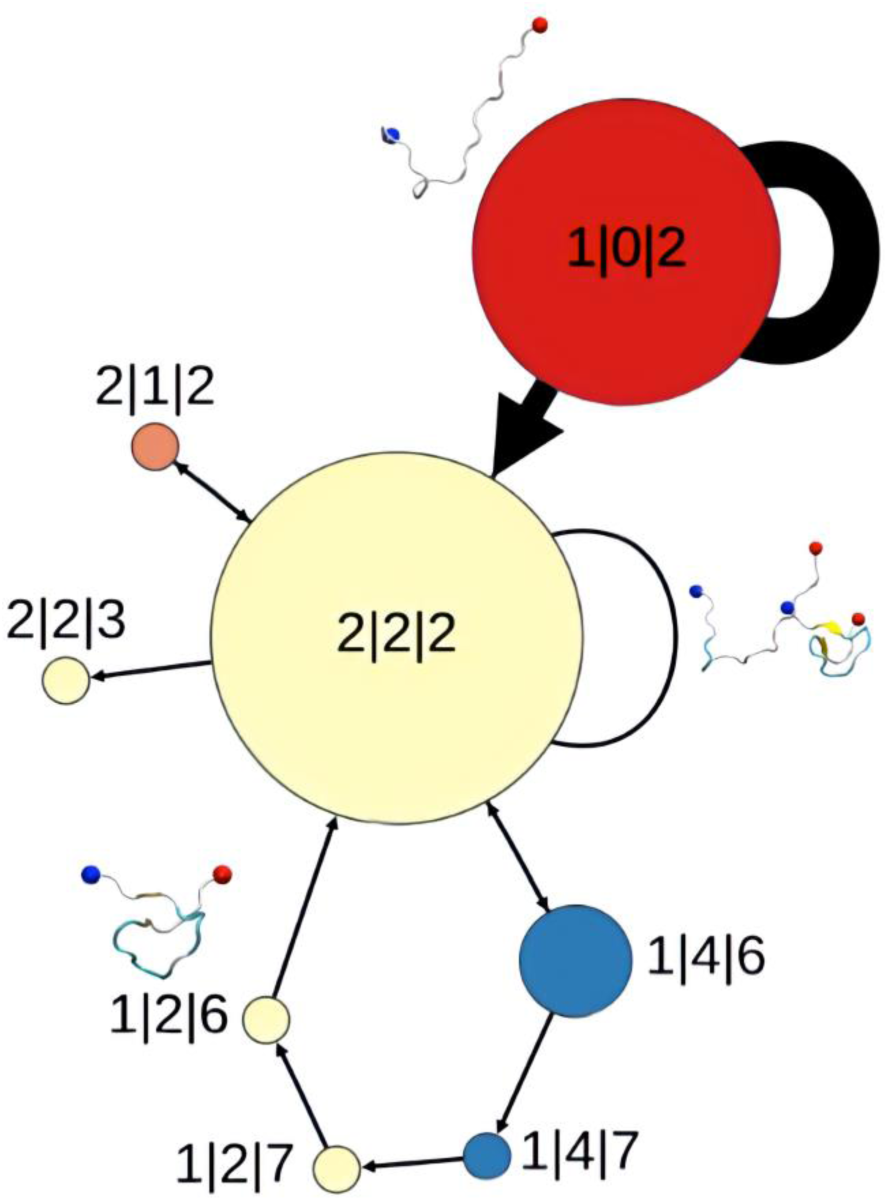
Transition network of dimer formation by two peptides with identical sequence of 25 amino acid residues. The size of the nodes is proportional to the state population and the thickness of the arrows is proportional to the number of transitions between the states. Next to each node the corresponding state attributes are given: oligomer size | number of amino acids in β- strand | compactness. For three of the state representative conformations are shown. The color of the nodes corresponds to the average number of amino acids in β-strand conformations: red = low, yellow = medium, blue = high β-strand content.

Fig. 8 describes the aggregation process where an elongated monomer (state 1|0|2) and a compact one (state 1|2|6 or state 1|4|6) come together and form an elongated dimer (state 2|2|2). The monomer states 1|2|7 and 1|4|7 do not directly assemble into a dimer but are first converted to state 1|2|6. The fluctuations in the β-strand content of the monomers can be clearly seen in the TN.

## 3 Summary

We provided step-by-step guides and necessary files for running MD simulations of peptide aggregation using GROMACS and analyzing these simulations in terms of oligomer size, inter-peptide contacts, Markov state models and transition networks. Peptide and protein aggregation is associated with a number of diseases, such as Alzheimer’s and Parkinson’s disease, and is thus under intensive study using both experimental and computational techniques [1,2]. For the latter, especially MD simulations with atomic resolution on the microsecond time scale have become an essential tool to investigate the relationship between sequence, conformational properties and aggregation of peptides or proteins [5]. As MD simulations produce a large amount of data, it is important to develop tools that extract information from the MD trajectories that provide key insight into the process under study. In terms of peptide/protein aggregation obvious key questions are whether oligomers formed during the simulation, how large these are, and by what interactions the aggregation process is driven. These questions can be answered by calculating the oligomer sizes and the inter-residue contacts within the oligomers, for which a Python script is provided here. To gain insight into the aggregation pathways, appropriate network models need to be deduced from the MD data. We presented two possibilities to calculate such network models, Markov state models and transition networks. The former are based on kinetic clustering of the MD data and thus elucidate key insight into the kinetically relevant aggregation pathways [11], as demonstrated here for the dimerization of the Aβ_16-22_ peptide. Transition networks (TNs), on the other hand, are based on conformational clustering and thus provide more structural details about the different oligomers that were sampled during the aggregation process. Moreover, the user can easily control how much detail should be presented in the TNs, which can reach from coarse-grained TNs with the oligomer size as the only descriptor to very fine-grained TNs with several descriptors per state to distinguish the oligomers of the same size from each other [12-16]. The experience from our lab shows that MSMs and TNs complement each other and it is thus advisable to calculate both kinds of networks from the MD data. It should be noted though that MSMs require converged MD data, which usually implies tens of microseconds of MD sampling, as otherwise they cannot be constructed.

In summary, the determination of oligomer sizes, contact maps, MSMs, and TNs are recommended for the analysis of MD trajectories studying peptide or protein aggregation. To realize such analysis the necessary files and explanations are provided in this chapter.

## Appendix

### Appendix A: Input files for the MD simulation

In the following, the five .*mdp* files required to follow the MD simulation protocols in section 2.1 are provided.

#### “ions.mdp” for adding ions

~~~
;; ions.mdp
; Run setup
integrator = steep
emtol = 1000
emstep = 0.01
nsteps = 2000
; Neighbor search
cutoff-scheme = Verlet
pbc = xyz
; Electrostatics and vdW
coulombtype = PME
pme-order = 4
fourierspacing = 0.1
rcoulomb = 1.2
rvdw = 1.2
~~~

#### “em.mdp” for energy minimization

~~~
;; em.mdp
; Run setup
integrator = steep
emtol = 500
emstep = 0.001
nsteps = 2000
nstxout = 100
; Neighbor searching
cutoff-scheme = Verlet
nstlist = 20
ns-type = grid
pbc = xyz
; Electrostatics
coulombtype = PME
pme-order = 4
fourierspacing = 0.1
rcoulomb = 1.2
; VdW
rvdw = 1.2
~~~

#### “nvt.mdp” for first equilibration in the NVT ensemble

~~~
;; nvt.mdp
define = -DPOSRES
; Run setup
integrator = md
dt = 0.002 ; 2 fs
nsteps = 50000
; Output control
nstxout = 5000
nstvout = 5000
nstfout = 5000
nstlog = 500
nstenergy = 500
; Bonds
constraints = all-bonds
constraint-algorithm = LINCS
lincs-order = 4
lincs-iter = 1
; Neighbor searching
cutoff-scheme = Verlet
nstlist = 20
ns-type = grid
pbc = xyz
; Electrostatics
coulombtype = PME
pme-order = 4
fourierspacing = 0.1
rcoulomb = 1.2
; VdW
rvdw = 1.2
; T coupling is on
tcoupl = v-rescale
tc-grps = Protein Non-Protein
tau-t = 0.1 0.1
ref-t = 300 300
nsttcouple = 10
; P coupling is off
pcoupl = no
; Velocity generation
gen-vel = yes
gen-temp = 300
continuation = no
~~~

#### “npt.mdp” for second equilibration in the NPT ensemble

~~~
;; npt.mdp
define = -DPOSRES
; Run setup
integrator = md
dt = 0.002
nsteps = 100000
; Output control
nstxout = 5000
nstvout = 5000
nstfout = 5000
nstlog = 500
nstenergy = 500
; Bonds
constraints = all-bonds
constraint-algorithm = LINCS
~~~

#### “md.mdp” for MD production run

~~~
;; md.mdp
; Run setup
integrator = md
dt = 0.002
nsteps = 500000000 ; 500 million
; Output control
nstxout = 0
nstvout = 0
nstfout = 0
nstlog = 2500
nstenergy = 2500
nstxout-compressed = 2500
compressed-x-grps = System
; Bonds
constraints = all-bonds
constraint-algorithm = LINCS
lincs-order = 4
; Neighbor searching
cutoff-scheme = Verlet
Nstlist = 20
ns-type = grid
pbc = xyz
; Electrostatics
coulombtype = PME
pme-order = 4
fourierspacing = 0.1
rcoulomb = 1.2
; VdW
rvdw = 1.2
; T coupling is on
tcoupl = v-rescale
tc-grps = Protein Non-Protein
tau-t = 0.1 0.1
ref-t = 300 300
nsttcouple = 10
; Pressure coupling is off
pcoupl = Parrinello-Rahman
pcoupltype = isotropic
tau_p = 2.0
ref_p = 1.0
compressibility = 4.5e-5
; Velocity generation
gen-vel = no
gen-temp = 300
continuation = yes
~~~

### Appendix B: Python script for the calculation of the oligomerization state and contact maps

~~~
import numpy as np
import copy as cp
import subprocess as sp
import os as os
import shutil as sh
import MDAnalysis as mdana
import sys
from MDAnalysis.analysis.distances import distance_array
import networkx as nx
import pandas as pd
import mdtraj as md
import matplotlib
matplotlib.use(“TkAgg”)
from matplotlib import pyplot as plt
#input parameters
ref_structure=sys.argv[1]
traj=sys.argv[2]
Min_Distance=int(sys.argv[3])
#structure parameters
topology = md.load(ref_structure).topology
trajectory = md.load(traj, top=ref_structure)
frames=trajectory.n_frames #Number of frames
chains=topology.n_chains #Number of chains
atoms=int(topology.n_atoms/chains) #Number of atoms in each monomer
AminoAcids = int(topology.n_residues/chains)-2 #Number of residues per chain
(‘-2’ to not count the N- and C- cap residues as individual residues)
isum=1
atoms_list=[]
atomsperAminoAcid=[]
residue_list=[]
for residue in topology.chain(0).residues:
 atoms_list.append(residue.n_atoms)
 residue_list.append(residue)
 ‘, ‘.join(map(lambda x: “‘” + x + “‘“, str(residue_list)))
#The N- and C- cap residues are part of the 1st and last residue index. If
no N- and C- cap residues for the protein, comment the line below using “#”
del residue_list[0]; del residue_list[-1]
for ii in range(len(atoms_list)):
 isum = isum + atoms_list[ii]
 atomsperAminoAcid.append(isum)
atomsperAminoAcid.insert(0, 1)
#The N- and C- cap residues are part of the 1st and last residue index. If
no N- and C- cap residues for the protein, comment the line below using “#”
del atomsperAminoAcid[1]; del atomsperAminoAcid[-2]
# Create Universe
uni = mdana.Universe(ref_structure,traj)
n,t = list(enumerate(uni.trajectory))[0]
box = t.dimensions[:6]
atom_Groups = [[] for x in range(chains)]
m_start=0
for m in range(0,chains):
 m_end = atoms * (m+1)
 atom_Groups[m].extend([uni.select_atoms(‘bynum ‘+ str(m_start) + ‘:’ +
str(m_end))])
 m_start = m_end + 1
fileout1 = open(‘oligomer-groups.dat’,’w’)
fileout2 = open(‘oligomer-states.dat’,’w’)
for tt in uni.trajectory[1:]:
 fileout1.write (str(tt.frame) + ‘\t’)
 fileout2.write (str(tt.frame) + ‘\t’)
 mySet = set([])
 graph = []
 atom_Groups = [[] for x in range(chains)]
 m_start=0
 for m in range(0,chains):
 m_end = atoms * (m+1)
 atom_Groups[m].extend([uni.select_atoms(‘bynum ‘+ str(m_start) + ‘:’
+ str(m_end))])
 m_start = m_end + 1
 count = 0
 for i in range(chains-1):
 for j in range(i+1,chains):
 dist = dis-
tance_array(atom_Groups[i][0].positions,atom_Groups[j][0].positions,box).min
()
 if(dist <= Min_Distance):
 my_tuple = (i,j)
 mySet.add(my_tuple)
 graph = nx.from_edgelist(mySet)
 for i in range(chains):
 if i not in list(graph):
 fileout1.write (‘[‘+ str(i)+’]’ + ‘\t’)
 fileout2.write (‘1’ + ‘\t’)
 else:
 fileout1.write (str(list(nx.node_connected_component(graph, i)))
+ ‘\t’)
 fileout2.write (str(len(list(nx.node_connected_component(graph,
i)))) + ‘\t’)
 fileout1.write (‘\n’)
 fileout2.write (‘\n’)
fileout1.close()
fileout2.close()
# Get oligomerization data
OligoStates = [[0 for z in range(chains)] for x in range(frames+1)]
file = open(“oligomer-groups.dat”,’r’)
line = file.readline()
j = 0
while line:
 temp = line.split(‘\t’)
 for k in range(chains):
 OligoStates[j][k] = temp[k + 1][1:-1].split(‘,’)
 j += 1
 line = file.readline()
file.close
# Create contact matrix
ContactMap = [[0 for x in range(AminoAcids)] for y in range(AminoAcids)]
# Create atom groups for each amino acid of each monomer
AtomGroups = [[] for x in range(chains)]
for m in range(0,chains):
 for aa in range(0,AminoAcids):
AtomGroups[m].extend([uni.select_atoms(‘bynum ‘+str(atoms*m + at-
omsperAminoAcid[aa])+’:’+str(atoms*m + atomsperAminoAcid[aa + 1] - 1))])
count = 0
for n,t in enumerate(uni.trajectory[1:]):
 on = 0
 Groups = []
 for i in OligoStates[n]:
 if len(i) > 1:
 on = 1
 Groups.extend([i])
 Set = set(tuple(x) for x in Groups)
 Groups = [list(x) for x in Set]
if on == 1:
# Calculate dimension of the box to considered PBC
 box = t.dimensions[:6]
 for g in Groups:
 # Calculate contacts
 for n1,i in enumerate(g):
 for j in g[(n1 + 1)::]:
 count += 1
 for n2,atoms1 in enumerate(AtomGroups[int(float(i))]):
 for n3,atoms2 in enumer-
ate(AtomGroups[int(float(j))]):
 if ((dis-
tance_array(atoms1.positions,atoms2.positions,box).min()) <= Min_Distance):
 ContactMap[n2][n3] +=1
 ContactMap[n3][n2] +=1
#print(count)
Norm_ContactMap = np.true_divide(ContactMap,float(count)*2.0)
# Save contact map in a file
fileout = open (‘contact-map.dat’,’w’)
for i in Norm_ContactMap:
 for j in i:
 fileout.write (str(j) + ‘\t’)
 fileout.write (‘\n’)
fileout.close()
#Highest Oligomer size in each frame
states=open(‘oligomer-states.dat’, ‘r’)
ter=states.readlines()[0:frames+1]
result=[]
for freq in (ter):
 result.append([int(hist) for hist in freq.strip().split(‘\t’)[1:]])
fileout3 = open (‘oligo-highest-size.dat’, ‘w’)
for oli_count in range(len(ter)):
fileout3.write(“{} {} {}\n”.format(oli_count, ‘\t’,
np.max(result[oli_count])))
fileout3.close()
# Block Average
size_data = np.loadtxt(‘oligomer-states.dat’)
window = 25 #over how many frames the running average is to be calculated weights = np.repeat(1.0,window)/window
size_data_m = np.convolve(size_data[:,1],weights,’valid’)
fileout4 = open(‘oligo-block-average.dat’, ‘w’)
for t,b in enumerate(size_data_m):
 fileout4.write(“{} {} {}\n”.format(t, ‘\t’, b))
fileout4.close()
~~~

### Appendix C: Tcl scripts for the transition network analysis

#### TNA.tcl

~~~
# Procedure - protein Compactness
proc NPMI {chs fr} {
 set sel [atomselect top “chain $chs” frame $fr]
 set eig [lsort -increasing -real [lindex [measure inertia $sel eigen-
vals] 2]]
 set shape [expr round([lindex $eig 0]/[lindex $eig 2]*10)]
 $sel delete
 return $shape
}
# Procedure - Amino acids in beta-sheet conformation
proc beta {chs fr} {
 set beta [atomselect top “chain $chs and name CA and (betasheet or
sheet or beta_sheet or extended_beta or bridge_beta)” frame $fr]
 set nb [expr [$beta num]]
 $beta delete
 return $nb
}
# Main code
# Read input files
set input1 [lindex $argv 0]
set input2 [lindex $argv 1]
# Load trajectory
mol new $input1
animate delete beg 0 end 0
animate read xtc $input2 waitfor all
# Select peptide length
set pep [lindex $argv 2]
set pepno [lindex $argv 3]
# Define chain id for each protein
set ch 0
set nAA [expr $pep*$pepno]
set TS {}
[atomselect top all] set chain 0
# Assign chain IDs
for {set i 0} {$i<$nAA} {incr i $pep} {
 [atomselect top “residue $i to [expr $i+$pep-1]”] set chain $ch
 incr ch
}
# Define cutoff
distance set cutoff 4
set d $cutoff
set oligomers {}
# Calculate the number of frames
set nf [molinfo top get numframes]
puts “nf = $nf”
# Open the transition matrix and states attributes files for writing
set fil1 [open “Transition-matrix.dat” w]
set fil2 [open “State-Attributes.csv” w]
set states {}
set prevolig {}
set S_unique {}
# Cycle through frames
for {set j 0} {$j<$nf} {incr j} {
 set cnt 0
 # Go to a specific frame
 puts “frame $j”
 animate goto $j
 display update
 mol ssrecalc top
 # Initialize some variables
 set oligomer {}
 set oldolig {} set olig {}
 set transition {}
 # Cycle through monomers
 for {set i 0} {$i<$pepno} {incr i} {
 # Check if the current peptide is already part of the current oligo-
mer
 if {[lsearch $oldolig $i] == −1} {
 set cnt {}
 lappend cnt $i
 # Identify neighboring chains within distance cutoff
 set res1 [atomselect top “(within $d of chain $i) and not chain
$i”]
 set length [llength [$res1 get chain]]
 set neighbor [lindex [lsort -unique [$res1 get chain]] 0]
 $res1 delete
 while {$length != 0} {
 lappend cnt $neighbor
 set res [atomselect top “(within $d of chain $cnt) and not
chain $cnt”]
 set length [llength [$res get chain]]
 set neighbor [lindex [lsort -unique [$res get chain]] 0]
 $res delete
 }
 set oldolig [concat $oldolig $cnt]
 # Update the list of peptides that belong to the same oligomer
 lappend olig $cnt
 # Update the list with the oligomer sizes
 lappend oligomer [llength $cnt]
 }
 }
 # Identify transitions
 if { $cnt >= 0 } {
 set n 1
 foreach o $olig {
 # Search for oligomer within the previous oligomer list
 set so [lsearch $prevolig $o]
 if {$so>=0} {
 lappend transition [list [lindex $prevolig $so] $o]
 } elseif {[llength $o] == 1} {
 # Search for monomers within previous oligomers (oligomer
broken into monomers)
 foreach p $prevolig {
 set so1 [lsearch $p $o]
 if {$so1>=0} {
 lappend transition [list $p $o]
 }
 }
 } elseif {[llength $o] > 1} {
 # If the oligomer is larger than 1 look for individual pep-
tides in the previous oligomer list (oligomer formed)
 foreach ol $o {
 set so2 [lsearch $prevolig $ol]
 if {$so2>=0} {
 lappend transition [list [lindex $prevolig $so2] $o]
 } else {
 # Search for individual peptides within previous oligo-
mers
 foreach pl $prevolig {
 set so3 [lsearch $pl $ol]
 if {$so3>=0} {
 lappend transition [list $pl $o]
 }
 }
 }
 }
 incr n
 }
 }
 set prevolig $olig
 set oldoligomer $oligomer
 set a2 {}
 # For each transition identify the aggregation states
 foreach t1 [lsort -unique $transition] {
 set a1 {}
 set b1 {}
 set S {}
 foreach t2 $t1 {
 set frame [expr $j-$f]
 animate goto $frame
 mol ssrecalc top
 lappend a1 [llength $t2]
 set ss {}
 set O [llength $t2]
 # set be 0
 #Monomer, call procedures
 if {$O == 1 } {
 set Sh [NPMI $t2 $frame]
 set be [beta $t2 $frame]
 } else {
 #Oligomer, call procedures
 set Sh [NPMI $t2 $frame]
 set be [beta $t2 $frame]
 }
 # Assign a state consisting of three order parameters
 set states [join $O|[expr round($be/$O)]|$Sh ““]
 lappend S $states
 }
 lappend a2 $a1
 set l [lindex $a1 0]
 set k [lindex $a1 1]
 if {$cnt!=0} {
 lappend TS $S
 }
 }
 }
}
# Write transition matrix and states attributes to files
if {1} {
 set S_unique [lsort -unique [join $TS]]
 set bbins [llength $S_unique]
 for {set i 0} {$i < $bbins} {incr i} {
 for {set j 0} {$j < $bbins} {incr j} {
 set b($i,$j) 0
 }
 }
 foreach trans $TS {
 set i [lsearch $S_unique [lindex $trans 0]]
 set j [lsearch $S_unique [lindex $trans 1]]
 set b($i,$j) [expr $b($i,$j)+1]
 }
 set row2 {}
 for {set i 0} {$i < $bbins} {incr i} {
 set row2 {}
 for {set j 0} {$j < $bbins} {incr j} {
 lappend row2 $b($i,$j)
 }
 puts $fil1 $row2
 puts $row2
 }
 # Create attributes file
 set all_states {}
 set count 0
 foreach v $TS {
 if {$count < $pepno} {
 lappend all_states [lindex $v 0]
 lappend all_states [lindex $v 1]
 } else {
 lappend all_states [lindex $v 1]
 }
 incr count
 }
 set id 1
 puts $fil2 “id state oligomer beta-sheet compactness population”
 foreach val $S_unique {
 puts $fil2 “$id $val [lindex [split $val ““] 0] [lindex [split
$val ““] 2] [lindex [split $val ““] 4] [llength [lsearch -all $all_states
$val]]”
 incr id
 }
}
close $fil1
close $fil2
exit
~~~

#### convert2csv.tcl: for converting the transition matrix to csv format

~~~
#!/usr/bin/tclsh
proc splitby { string spl_str } {
 set lst [split $string $spl_str]
 for { set cnt 0 } { $cnt < [llength $lst] } { incr cnt } {
 if { [lindex $lst $cnt] == “” } {
 set lst [lreplace $lst $cnt $cnt]
 incr cnt −1
 }
 }
 return $lst
}
set input [lindex $argv 0]
set output1 [lindex $argv 1]
set fil1 [open $input r]
set fil2 [open $output1 w]
array unset a
set i 1
set firstR {}
# Read input matrix into variable “a”
while {[gets $fil1 line1] >=0} {
 set firstR [concat $firstR “;$i”]
 set row [splitby $line1” “]
 set j 1
 foreach r $row {
 set a($i,$j) $r
 incr j
 }
 incr i
}
set n [expr $i-1]
# Write output matrix to file
puts $fil2 $firstR
for {set i 1} {$i <= $n} {incr i} {
 set var $i
 for {set j 1} {$j <= $n} {incr j} {
 set var [concat $var “;$a($i,$j)”]
 }
 puts $fil2 $var
}
close $fil1
close $fil2
quit
~~~

